# Species composition determines bioplastics production in photosynthetic microbiomes: strategy to enrich cyanobacteria PHB-producers

**DOI:** 10.1101/2023.05.30.542808

**Authors:** Beatriz Altamira-Algarra, Artai Lage, Joan García, Eva Gonzalez-Flo

**Author notes:** Corresponding author. GEMMA-Group of Environmental Engineering and Microbiology. Department of Civil and Environmental Engineering. Escola d’Enginyeria de Barcelona Est (EEBE). Universitat Politècnica de Catalunya-BarcelonaTech. Av. Eduard Maristany 16. Building C5.1. E-08019 Barcelona. Spain. **Funding** This research has received funding from the European Union’s Horizon 2020 research and innovation programme under the grant agreement No 101000733 (project PROMICON). B. Altamira-Algarra thanks the Agency for Management of University and Research (AGAUR) for her grant [FIAGAUR_2021]. E. Gonzalez-Flo would like to thank the European Union-Next Generation EU, Ministry of Universities and Recovery, Transformation and Resilience Plan for her research grant [2021UPF-MS-12].

## Abstract

The aim of this study was to set the operating mode in regards to nutrients, temperature and light to use as a strategy to enrich a microbiome rich in cyanobacteria in polyhidroxybutyrate (PHB)-producers in order to enhance this biopolymer production. Alternate growth and accumulation phases were conducted for 179 days in a 3 L photobioreactor. Although, presence of green microalgae potentially reduced PHB production, the microbiome produced up to 22 % dry cell weight (dcw) PHB. Results suggested that this methodology could be applied to a robust microbiome rich in cyanobacteria to boost PHB production.

## 1. Introduction

Polyhydroxybutyrate (PHB) is a biopolymer synthetized by numerous bacterial species as intracellular carbon and energy storage being fully biodegradable into CO_2_ and H_2_O (Jendrossek and Handrick, 2002). Due to their physicochemical properties, PHB can be used in different fields, such as in packaging or medicine (Manikandan et al., 2020; Mohammadalipour et al., 2023). Industrial PHB production is nowadays based exclusively on “pure” microbial cultures, which raises production expenses and makes it difficult for PHB to be price competitive with traditional plastics (Tan et al., 2021).

Use of microbial “mixed” cultures (microbiomes) for PHB production has the potential to comparatively reduce operational costs of pure cultures since they could be operated in open systems not requiring sterilization, and utilize cheap by-products and waste streams as feed (Fradinho et al., 2013a; Mohamad Fauzi et al., 2019; Reis et al., 2011). This approach would engage the integration of the biopolymer production process within the circular economy concept. To increase biopolymer production in microbiomes two complementary strategies can be followed: (i) optimizing the culture to enhance the presence of PHB-producers and (ii) modifying operating parameters with an effect on PHB metabolic routes and subsequent synthesis. The called feast and famine (FF) strategy has been successfully applied to enrich cultures in PHB-storing organisms (Oliveira et al., 2017; Sagastume et al., 2017; Sruamsiri et al., 2020). FF regime consists in a transient period of carbon source availability in which microorganisms store PHB (feat phase). Followed by a prolonged period without carbon addition (famine phase), where microorganisms use the stored biopolymer as an internal carbon source. Repeated FF cycles create a selection pressure favorable for microorganisms with the capacity to store PHB. Productivities up to 90 % dry cell weight (%dcw) of PHB have been produced by heterotrophic microbiomes at laboratory scale, which is comparable to those obtained by pure cultures (Johnson et al., 2009; Serafim et al., 2004). However, until now, most of the experiments on PHB production using FF have been performed by heterotrophic microorganisms, like activated sludge using volatile fatty acids as substrate (Crognale et al., 2022; Estévez-Alonso et al., 2022; Johnson et al., 2009; Serafim et al., 2004), while there is a wide diversity of bacterial species that can produce this biopolymer and numerous operation conditions to be tested.

Photosynthetic microorganisms, such as cyanobacteria, are also able to produce PHB. Although production rates are lower than those obtained by heterotrophic bacteria (Monshupanee and Incharoensakdi, 2014; Rueda et al., 2020a; Sharma and Mallick, 2005a), the interest of using a photosynthetic culture is the feasibility of using CO_2_ and sunlight for biomass growth and biopolymer synthesis (Rueda et al., 2020a). Laboratory research on PHB production by cyanobacteria has been done using monocultures of, for example, *Nostoc* sp., *Synechocystis* sp. and *Synechococcus* sp. (Ansari and Fatma, 2016; Monshupanee and Incharoensakdi, 2014; Rueda et al., 2022a, 2020b; Sharma and Mallick, 2005b). These studies examined the effect of different operation factors on PHB production, such as culture conditions (*e.g.* photoautotrophic, mixotrophic or photoheterotrophic). Results showed that under photoheterotrophic regime, PHB production is enhanced reaching values up to 30 %dcw (Rueda et al., 2022a), making the process a possible candidate for industrial application.

A mixed culture enriched with cyanobacteria would in effect combine the aforementioned advantages of working with microbiomes and cyanobacteria, potentially overcoming the production rates obtained by cyanobacteria monocultures up to now. Nevertheless, to the authors’ knowledge, photosynthetic microbiomes enriched with cyanobacteria have only been tested for PHB production in (Altamira-Algarra et al., 2022; Arias et al., 2018; Rueda et al., 2020c), obtaining a PHB production up to 14 %dcw PHB (Altamira-Algarra et al., 2022). In (Altamira-Algarra et al., 2022) the effect of the number of days under light and the presence of inorganic and organic carbon on PHB production by a photosynthetic microbiome was evaluated. Outcomes revealed that the addition of an organic carbon source (acetate) greatly triggered biopolymer synthesis and reactors could be under dark during PHB production. the presence of inorganic carbon (as bicarbonate) had no notable impact on biopolymer synthesis. These findings were used to establish the operating procedure in a next step using reactors with higher volumes, as well as, to evaluate the feasibility of increasing PHB production by enhancing the presence of organisms producing the biopolymer.

In addition, one research gap that needs to be addressed is how to maintain productive cyanobacterial cultures for the long-term generation of bioproduct. Unfortunately, most research undertaken to date has been limited to small-scale experiments with a short time frame, typically lasting only a few weeks. There have been no attempts to maintain cultures in the long term (on the order of several months) for continuous bioproduct generation, which is of important value for industrialization of bioproduct synthesis. Thus, in this work, a photosynthetic microbiome was cultivated in a photobioreactor with a sequential operation for a total of 179 days. The study evaluated a novel methodology based on the FF strategy to enhance with PHB-producers a microbiome rich in cyanobacteria.

## 2. Material and methods

### 2.1. Inoculum and experimental set-up

Microbiome named CC, isolated in (Altamira-Algarra et al., 2022) was used as the inoculum in 3 L glass cylindrical photobioreactors (PBRs) of 2.5 L working volume (Fig. A.1). This microbiome sample was collected from the Canal dels Canyars outlet (Gavà, Spain, 41°15’55.9“N 2°00’39.7”E), very near to the sea, and it was rich in the unicellular cyanobacteria *Synechococcus* sp. and the filamentous cyanobacteria *Leptolyngbya* sp. (Fig. A2). Illumination in reactors was kept at 30 klx (approx. 420 µmol·m^-2^·s^-1^) by 200W LED floodlight, placed at 15 cm from the reactors surface in 15:9 h light:dark cycles. pH was measured online with a pH probe (HI1001, HANNA instruments, Italy) placed inside the reactors and was controlled at around 7.5 (during growth phase) by a pH controller (HI 8711, HANNA instruments, Italy) activating an electrovalve which injected CO_2_ inside the reactors when pH reached 8.5. The pH data was saved in a 5 min interval in a computer with the software PC400 (Campbell Scientific). In the PHB-accumulation phases the pH was measured but not controlled in order to avoid IC presence. Reactors were continuously agitated by a magnetic stirrer ensuring a complete mixing and culture temperature was kept at 30 °C or 35 °C according to the test conditions.

### 2.2. Experimental strategy

A novel procedure based on the FF strategy used in PHB production by heterotrophic cultures was applied to the microbiome CC. PBRs were operated for 179 days during which growth/starvation phases were constantly conducted with the aim to enrich the microbiome with PHB-producers’microorganisms.

The PBRs were operated in semi-continuous regime. In a first conditioning period, cultures were grown by adding bicarbonate (as inorganic carbon, IC) to the medium, and when nitrogen (N) was depleted, a first PHB-accumulation phase started. Secondly, three cycles consisting of (i) growth and (ii) starvation phases were done to determine the optimal operating conditions to favour cyanobacteria growth together with PHB production. For that, the effect of three ecological stresses (i) N concentration; (ii) temperature and (iii) light were also evaluated. Each parameter was tested individually in one cycle. Finally, after establishing the best results for each parameter, 9 iterated repetitions of (i) growth and (ii) starvation phases were conducted to enrich the biome in PHB-producers. Note that only in the beginning of the first growth phase (in conditioning period) bicarbonate was supplemented to the medium.

Experiment was conducted as follows (Fig. 1, and for its description see the text below):

**Figure 1.**
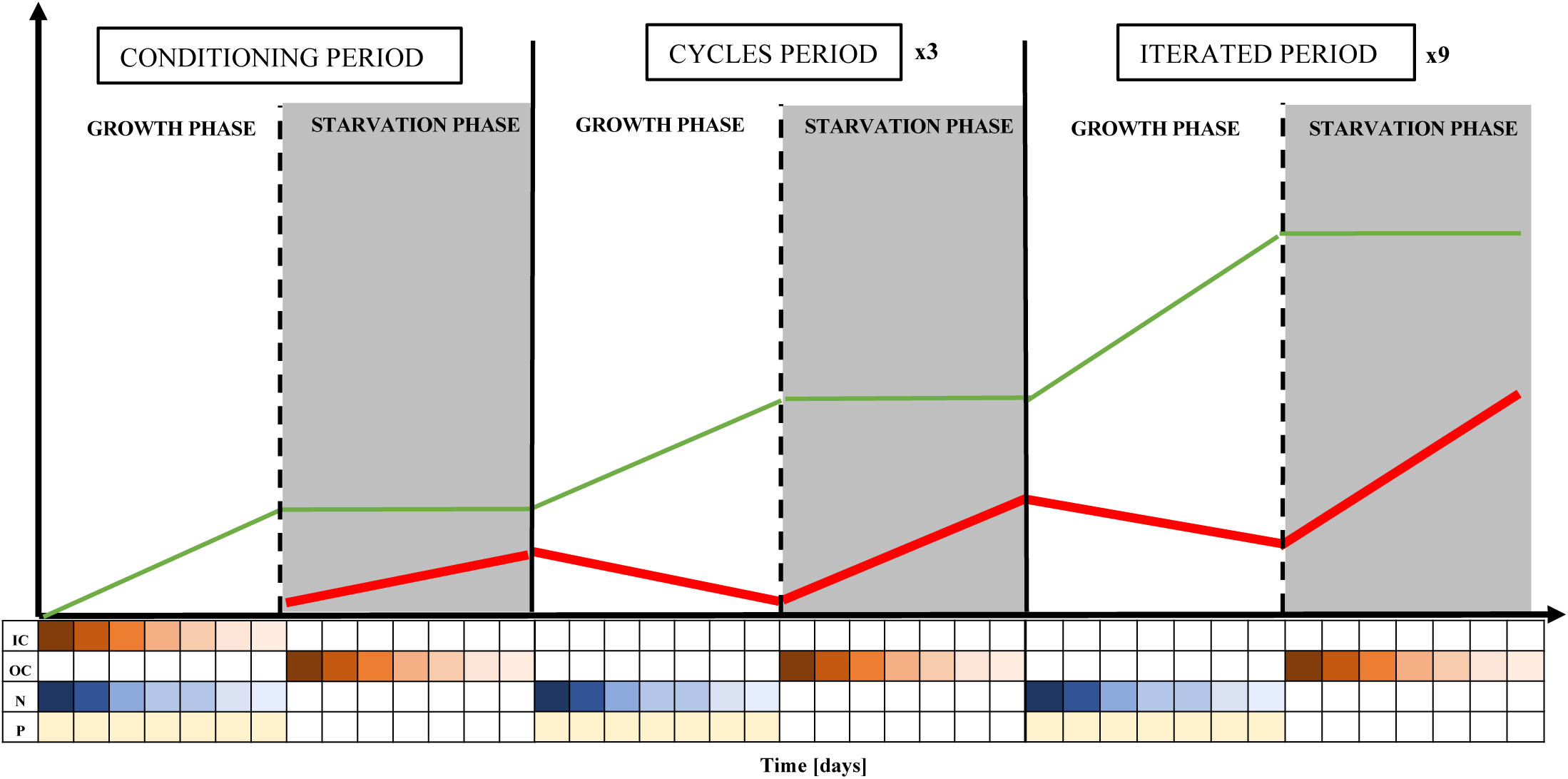
Schematic representation of the PBR semi-continuous operation applied for microbiome optimization for PHB-producing cyanobacteria populations. In green line is showed the assumed increase in PHB-producers biomass; in red, the assumed PHB evolution over time. Colours below the graph represent the concentration of the main nutrients in the experimental phases. The lighter the colour, the lower the concentration of the compound. Brown colour represents the inorganic carbon (IC) and the organic carbon (OC) concentration; blue colour represents the N and yellow is for P. White and grey colour in the figure indicates growth phase and starvation phase, respectively, in all the figures from this paper.

i. Conditioning period. Two PBRs were used here as duplicates:

- Growth phase: the PBRs were inoculated with 100 mg biomass (as volatile suspended solids, VSS)·L^-1^. BG-11 with modified concentrations of inorganic carbon (IC), N and P (100 mgIC·L^-1^ (expressed as C), 50 mgN·L^-1^ and 1 mgP·L^-^ was used as media for culture growth.
- Starvation phase: this step began when N was depleted. 600 mg acetate (Ac)·L^-1^ was added at this point and PBRs were enclosed with PVC tubes to keep the reactors under dark conditions. This first starvation phase was kept for 14 days.
ii. Cycles period.

Three cycles of (i) growth and (ii) starvation phases where conducted with 4 PBRs (duplicates for each parameter tested).

#### Cycle 1: Evaluation of N concentration during the growth phase

- Growth phase: First, biomass from the two PBR of the prior phase was mixed and divided based in volume in four PBRs. Initial biomass concentration was set at approximately 400 mgVSS·L^-1^ in the PBRs. Biomass was cultured with new BG-11 medium with modified concentrations of N and P, and without IC. N concentration was set at 25 mgN·L^-1^ for two PBRs, while the other two were inoculated with medium to have 50 mgN·L^-1^. P concentration was set at 0.1 mg·L^-1^ due to the sudden growth of green algae during the conditioning revealed by microscope observations. P was daily added to the medium to maintain 0.1 mg·L^-1^ in the PBRs (Table 1). The P was maintained during the whole growth phase by daily dosing a P solution of KH_2_PO_4_ to the PBRs.
- Starvation phase: The same operation mode was applied as in the starvation phase from the conditioning period. This phase and the coming starvation phases lasted 7 days.

**Table 1.** Stablished conditions used in each period of the experiment.

| Period | Phase | N (mg·L<sup>-1</sup>) | P (mg·L<sup>-1</sup>) | Ac (mg·L<sup>-1</sup>) | Lightness<br>(h of light:dark ) | T (°C) | Number<br>of PBR |
| --- | --- | --- | --- | --- | --- | --- | --- |
| Conditioning | Growth | 50 | 1 | - | 15:09 | 30 | 2 |
|  | Starvation | - | - | 600 | 0 | 30 | 2 |
| Cycle 1 | Growth | 25 (PBR 1 & 2)<br>50 (PBR 1 & 2) | 0.1 | - | 15:09 | 30 | 4 |
|  | Starvation | - | - | 600 | 0 | 35 | 4 |
| Cycle 2 | Growth | 25 | 0.1 | - | 15:09 | 30 | 4 |
|  | Starvation | - | - | 600 | 0 | 35 (PBR 1 & 2)<br>30 (PBR 3 & 4) | 4 |
| Cycle 3 | Growth | 25 | 0.1 | - | 15:09 | 30 | 4 |
|  | Starvation | - | - | 600 | 0 (PBR 1 & 2)<br>continuous (PBR 3 & 4) | 35 | 4 |
| Iterated period | Growth | 25 | 0.1 | - | 15:09 | 30 | 2 |
|  | Starvation | - | - | 600 | 0 | 35 | 2 |

#### Cycle 2: Evaluation of temperature during the starvation phase

- Growth phase: A certain volume of the PBRs usually ranging from 800 mL to 1,200 mL was discarded to purge the culture broth. The removed volume was calculated to set 400 mgVSS·L^-1^ as initial biomass concentration in the PBRs (similar to cycle 1). Discarded volume was replaced with new BG-11 medium with modified concentrations of N and P, and without IC (as bicarbonate). N concentration was set at 25 mgN·L^-1^ (best result from the prior cycle) and P, at 0.1 mg·L^-1^

- Starvation phase: Two reactors were set at 30 °C and the other two, at 35 °C. The other parameters were as in the conditioning phase.

#### Cycle 3: Evaluation of light during the starvation phase

- Growth phase: Equal operation mode as in cycle 2.
- Starvation phase: Two reactors were enclosed with PVC tubes for dark conditions while the other two were maintained in light:dark cycles as described before. Temperature was maintained at 35°C (best result from prior cycle).

iii. Iterated period.

A total of nine growth and starvation phases were conducted with the optimal values for the evaluated factors (N, temperature and light) obtained from the cycles period (Table). Biomass from PBR 1 & 2 from the prior phase was mixed and divided in two PBRs, while biomass from PBR 3 & 4 was discarded, thus 2 PBR were used in these cycles. A total of 9 iterated cycles of (i) growth and (ii) starvation phase were conducted. At the beginning of each growth phase, volume ranging from 800 mL to 1,200 mL was discarded to purge the culture broth and set 400 mgVSS·L^-1^ as initial biomass concentration in the PBRs (as in the previous cycles). Discarded volume was replaced with new BG-11 medium with modified concentrations of N and P, and without IC. Two PBRs were used as replica.

### 2.3. Analytical methods

At selected times, a 50 mL sample was taken from each PBR for analysis. Biomass concentration was determined by analysis of total suspended solids (TSS) and VSS as described in (Amerian Public Health Association, 2012). Turbidity was measured with turbidimeter (HI93703, HANNA Instruments). For a quick estimation of biomass content, VSS and turbidity were correlated by calibration curve (Fig. A.3):

To determine dissolved species, samples taken out from the reactors were previously filtered through a 0.7 μm pore glass microfiber filter. Nutrients (N and P) were measured as nitrate (N-NO_3_ ^-^) and phosphate (P-PO_4_ ^3-^) following Standard Methods (Amerian Public Health Association, 2012). Note that in BG-11 the only source of N is nitrate. The filtered sample was passed through a 0.45 μm pore size filter a second time to determine Ac by ion chromatography (CS-1000, Dionex Corporation, USA).

Biomass composition was monitored by bright light and fluorescence microscope observations (Eclipse E200, Nikon, Japan). Cyanobacteria and green algae were identified and classified following morphological descriptions (Komárek et al., 2020, 2011). Cell counting was done in a Neubauer chamber at the end of each starvation phase. Individual cells were counted until reach >400 cells to have a standard error lower than 5 % (Margalef, 1984).

### 2.4. PHB extraction and quantification

PHB analysis was adapted from methodology described in (Lanham et al., 2013). Briefly, 50 mL of mixed liquor were collected and centrifuged (4,200 rpm, 7.5 min), frozen at −80 °C overnight in an ultra-freezer (Arctiko, Denmark) and finally freeze-dried for 24 h in a freeze dryer (−110 °C, 0.05 hPa) (Scanvac, Denmark). 3-3.5 mg of freeze-dried biomass were mixed with 1 mL CH3OH with H2SO4 (20% v/v) and 1 mL CHCl_3_ containing 0.05 % w/w benzoic acid. Samples were heated for 5 h at 100 °C in a dry-heat thermo-block (Selecta, Spain). Then, they were placed in a cold-water bath for 30 min to ensure they were cooled. After that, 1 mL of deionized water was added to the tubes and they were vortexed for 1 min. CHCl_3_ phase, containing PHB dissolved, was recovered with a glass pipette and introduced in a chromatography vial containing molecular sieves. Samples were analysed by gas chromatography (GC) (7820A, Agilent Technologies, USA) using a DB-WAX 125-7062 column (Agilent, USA). Helium was used as the gas carrier (4.5 mL·min^-1^). Injector had a split ratio of 5:1 and a temperature of 230 °C. FID had a temperature of 300 °C. A standard curve of the co-polymer PHB-HV was used to quantify the PHB content.

### 2.5. Calculations

Total biovolumes (BV) in mm^3^·L-^1^ of each species (cyanobacteria (*Synechococcus* sp.) and the green microalgae) were calculated using the formula:

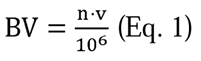

where n is the number of cells counted in a sample (cells·L^-1^) and v is the volume of each cell (μm^3^). 10^6^ is the unit conversion from µm^3^·mL to mm^3^·L^-1^. Cell volume was calculated by volumetric equations of geometric shape closest to cell shape. Biovolume of *Synechococcus* sp. was calculated by the volume equation of a cylinder and biovolume of green algae was obtained by the volume equation of an ellipsoid. Cell dimensions (length and width) were obtained from images of microscope observations (software NIS-Element viewer®) (Table A1).

Kinetic coefficients were calculated as follows:

Specific growth rate (d^-1^) was calculated using the general formula

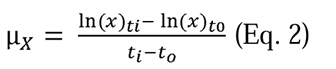

where ln(X)_ti_ and ln(X)_tn_ are the natural logarithms of the biomass concentration (mgVSS·L^-1^) at experimental day (t_i_) and at the beginning of the phase (t_0_), respectively. The terms t_i_ and at t_0_ are the time span (in days) at which µ_X_was calculated (when biomass concentration reached stationary phase).

Biomass volumetric production rate (mg·L^-1^·d^-1^) was calculated as:

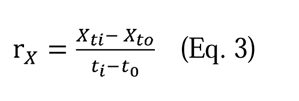

where X_ti_ (mg·L^-1^) and X_t0_ (mg·L^-1^) are the biomass concentration (in mgVSS·L^-1^) at time t_i_ (experimental day, when cell growth reached stationary phase) and at the beginning of the growth phase (t_0_). *i* is the total number of days that the growth phase lasted.

The nutrients (nitrogen) to biomass yield was calculated by (only during the growth phase):

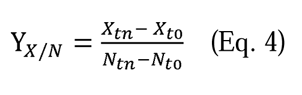

where X_tn_ (mg·L^-1^) and X_t0_ (mg·L^-1^) are the biomass concentration (in mgVSS·L^-1^) at the end (t_n_) and at the beginning of the phase (t_0_). N_tn_ (mg·L^-1^) and N_t0_ (mg·L^-1^) are the nitrogen concentration (N-NO_3_^-^) at the end and at the beginning of each growth phase, respectively.

The specific consumption rate of nitrogen (mgN·mgVSS^-1^·d^-1^) was calculated as:

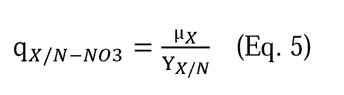

PHB volumetric production rate (C_PHB_ (mgPHB·L^-1^·d^-1^)) was obtained by:

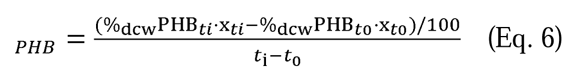

where %_dcw_PHB_tn_ and %_dcw_PHB_ti_ are the percentage of PHB respect biomass quantified at time *i* (experimental day) and at the beginning of the accumulation phase (t_0_). X and X are the biomass concentration (in mgVSS·L^-1^) at time *i* (experimental day) and at the beginning of the accumulation phase (t_0_), respectively.

The PHB yield on acetate (Ac) (Y_PHB/Ac_) was calculated on a COD-basis by:

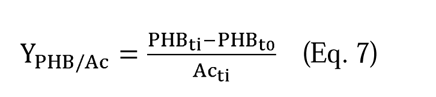

The amount of PHB produced (given as chemical oxygen demand (COD): 1.67 gCOD·gPHB^-1^) was obtained by multiplying the %dcw PHB produced per biomass concentration (in mgVSS·L^-1^) at time *i* (experimental day) and at the beginning (t_0_) of the accumulation phase. Ac_ti_ (mg·L^-1^) is the acetate concentration (given 1.07 gCOD·gAc^-1^) at the experimental day (t_i_) of the starvation phase. Ac_ti_ was calculated by subtracting the amount added (600 mgAc·L^-1^) from the amount of acetate left in the medium.

## 3. Results and discussion

### 3.1. Conditioning period

A first growth phase with 100 mgIC·L^-1^ (added as bicarbonate), 50 mgN·L^-1^ and 1 mgP·L^-1^ to reach a clearly detectable growth and biomass concentration was conducted. The average specific growth rate was 0.31 d^-1^ (Table 2) and after 7 days the microbiome grew up to 1,120 mgVSS·L^-1^ (Fig. 2A and Table 2). Nitrogen was assimilated during this growth phase at a specific consumption rate of 15 mgN·gVSS^-1^·d^-1^ (Table 2). These values were in accordance with previous work with monocultures of cyanobacteria under similar culture conditions (Rueda et al., 2022a, 2022b).

**Figure 2.**
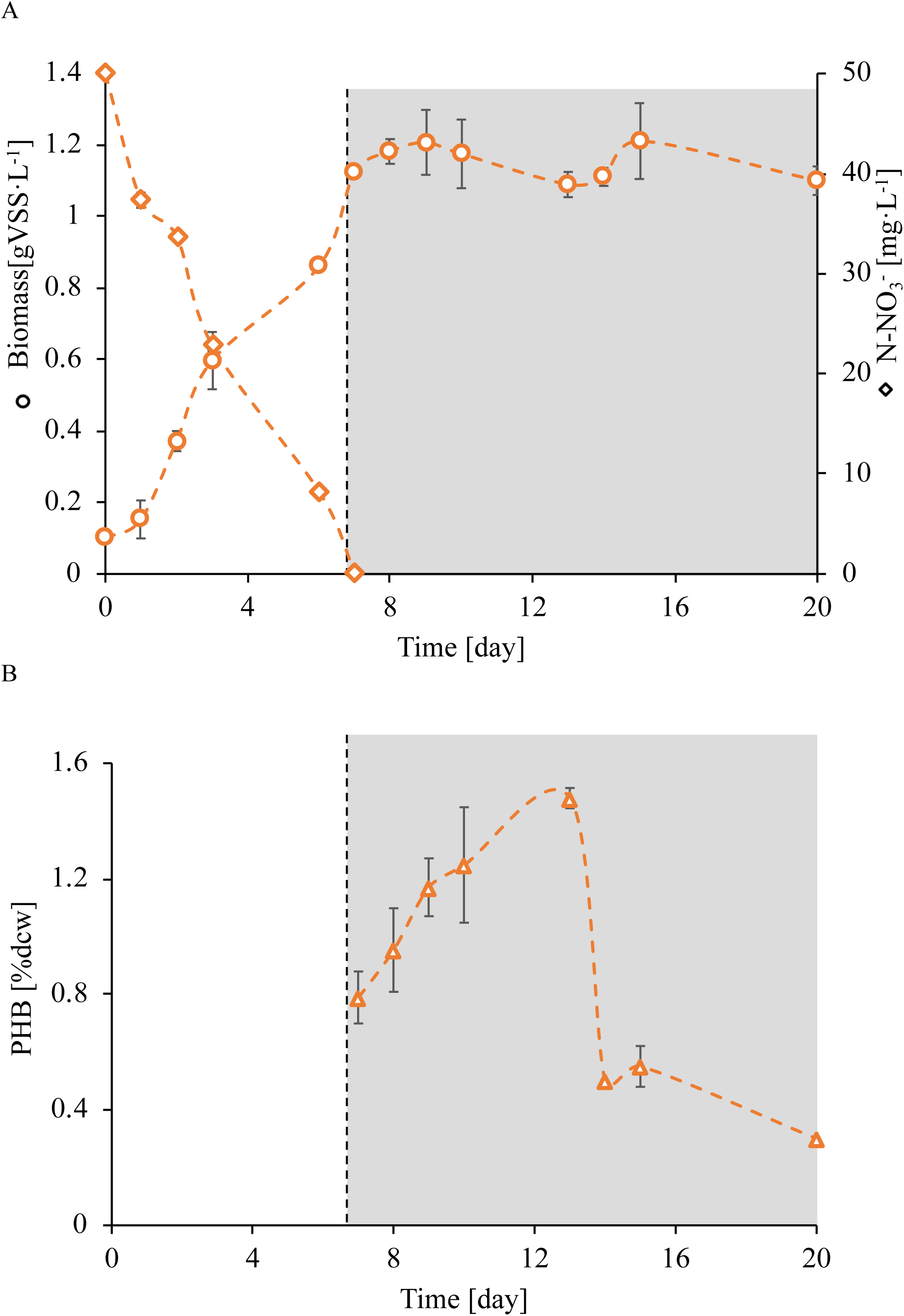
(A) Concentration changes in biomass (as volatile suspended solids, VSS) and nitrogen (as N-NO_3_) in PBR 1 & 2. (B) PHB production in PBR 1 & 2. Error bars indicate the standard deviation of the replicates. Nitrate was not measured during starvation phase. PHB was not measured in growth phase. Values of biomass (as VSS) were obtained following (Amerian Public Health Association, 2012).

**Table 2.** Average of the kinetic and stoichiometric parameters obtained during growth and accumulation phase of the conditioning period.

| Growth phase |  |
| --- | --- |
| Parameter | PBR 1 & 2 |
| VSS (mg·L<sup>-1</sup>) * | 1,120 |
| μ (d<sup>-1</sup>) * | 0.31 |
| □<sub>biomass</sub> (mgVSS·L<sup>-1</sup>·d<sup>-1</sup>) | 92.92 |
| q<sub>N</sub> (mgN·gVSS<sup>-1</sup>·d<sup>-1</sup>) | 15.35 |
| Y<sub>X/N</sub> | 20.45 |
| Accumulation phase |  |
| Parameter <sup>a</sup> | PBR 1 & 2 |
| PHB (%dcw) | 1.47 ± 0.04 |
| □<sub>PHB</sub> (mgPHB·L<sup>-1</sup>·d<sup>-1</sup>) | 1.29 ± 0.51 |

<sup>a</sup>Results at day seventh of the phase.

After seven days of reactor operation, N was depleted (Fig. 2A) and 600 mgAc·L^-1^ were added to the medium. This first starvation phase was maintained 14 days in order to follow PHB synthesis (Fig. 2B). Biomass concentration remained constant during this period (Fig. 2A). Regarding to biopolymer synthesis, it increased until day 7 in starvation, when 1.5 %dcw PHB was produced in both PBRs. However, from that day on, biopolymer synthesis decreased, and after 14 days in starvation, there was almost no PHB present; probably due to depletion of Ac and the consumption of the produced PHB. Consequence of this result, subsequent starvation phases lasted one week to avoid PHB consumption under dark, which would not favour growth of photosynthetic PHB-producers. Note that in this period the PHB synthesis was comparatively low.

The two PBRs were inoculated with a microbiome rich in the unicellular cyanobacteria *Synechococcus* sp. and the filamentous cyanobacteria *Leptolyngbya* sp. Unfortunately, a sudden growth of green algae was observed during this conditioning period (Fig. A.2). In fact, BV calculation disclosed that 80 % of the culture was composed by these microalgae. Consequently, to avoid green microalgae and favour cyanobacteria growth, P concentration at the beginning of each growth phase was fixed at 0.1 mg·L^-1^. Note that the biomass used as inoculum for the PBRs had been kept in modified BG-11 medium with a P concentration of 0.5 mg·L^-1^ (Altamira-Algarra et al., 2022). The P concentration set at the beginning of the conditioning period was 1 mgP·L^-1^ to sustain cell growth; however, we assumed that this value was too high and P stopped being growth-limiting for green algae.

### 3.2. Growth and PHB production under ecological stress

The conditioning period was done to obtain sufficient biomass concentration in the PBRs and initiate a first PHB accumulation step, which we assumed it would serve as the internal biopolymer for cell growth on the coming growth phase. After fourteen days in starvation (accumulation phase of the conditioning phase), biomass was equally divided into four PBRs and three cycles of (i) growth and (ii) starvation were conducted. In these cycles, the effect of N concentration at the beginning of the growth phase (cycle 1), temperature (cycle 2) and light (cycle 3) at the starvation phase was assessed.

To do so, no IC (as bicarbonate) was added at the growth phase to create a selection pressure favourable for microorganisms with the capacity to store PHB, because they would use the accumulated biopolymer during the starvation phase as an internal carbon source (Reis et al., 2011). However, CO_2_ was injected in the PBRs to control pH due to photosynthetic activity. Note that this CO_2_ presence, even small, is beneficial for biopolymer degradation, as cells need to resume photosynthetic activity in order to use the stored PHB (Klotz et al., 2016).

To support the growth on internal PHB, nutrients are required; thus, two N concentrations were evaluated (cycle 1). In two PBRs (PBR 1 and PBR 2) N concentration was lowered to 25 mgN·L^-1^ and in the other two PBRs (PBR 3 and PBR 4), N concentration was 50 mgN·L^-1^, the same as in the first growth phase (conditioning period). Biomass increased from 400 mgVSS·L^-1^ to 670 ± 100 mgVSS·L^-1^ in PBR 1 & 2 in seven days, and up to 1,300 ± 140 mgVSS·L^-1^ in PBR 3 & 4 in nine days (Fig. 3A and Table 2). The increase in biomass and the calculated kinetic parameters (Table 2 and Table 3) were higher in the latter PBRs because initial N concentration was also major (50 mgN·L^-1^). However, PHB production and productivity were lower (Table 3). In addition, by lowering N concentration at the beginning of the growth phase, this phase could be shortened to seven days; thus, reducing the days of the whole process, as 700 ± 100 mgVSS·L^-1^ was considered to be enough biomass concentration to start the accumulation step for next experiments. Therefore, N concentration was set at 25 mg·L^-1^at the beginning of each growth phase, and it lasted 7 days.

**Figure 3.**
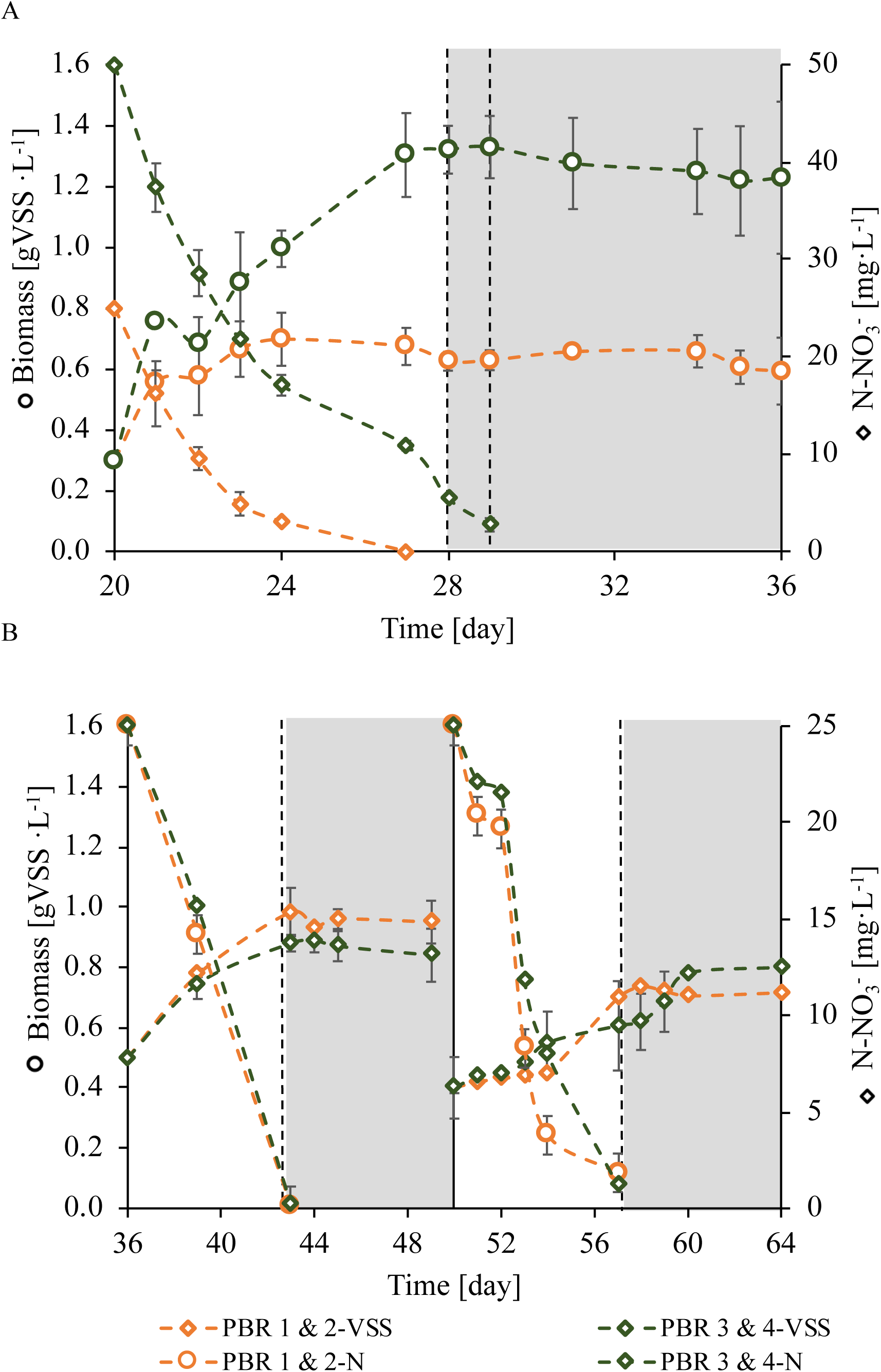
Concentration changes in biomass (as volatile suspended solids, VSS) and nitrogen (as N-NO_3_) in PBR 1 & 2 (orange) and PBR 3 & 4 (green) (A) during cycle 1 and (B) during cycle 2 and 3. First dashed line in (A) indicates the beginning of starvation phase in PBR 1 & 2; second dashed line stands for PBR 3 & 4. Continuous black line in (B) illustrates end of cycle 2 and beginning of cycle 3. Values of biomass (as VSS) were obtained by turbidity measurements.

**Table 3.**
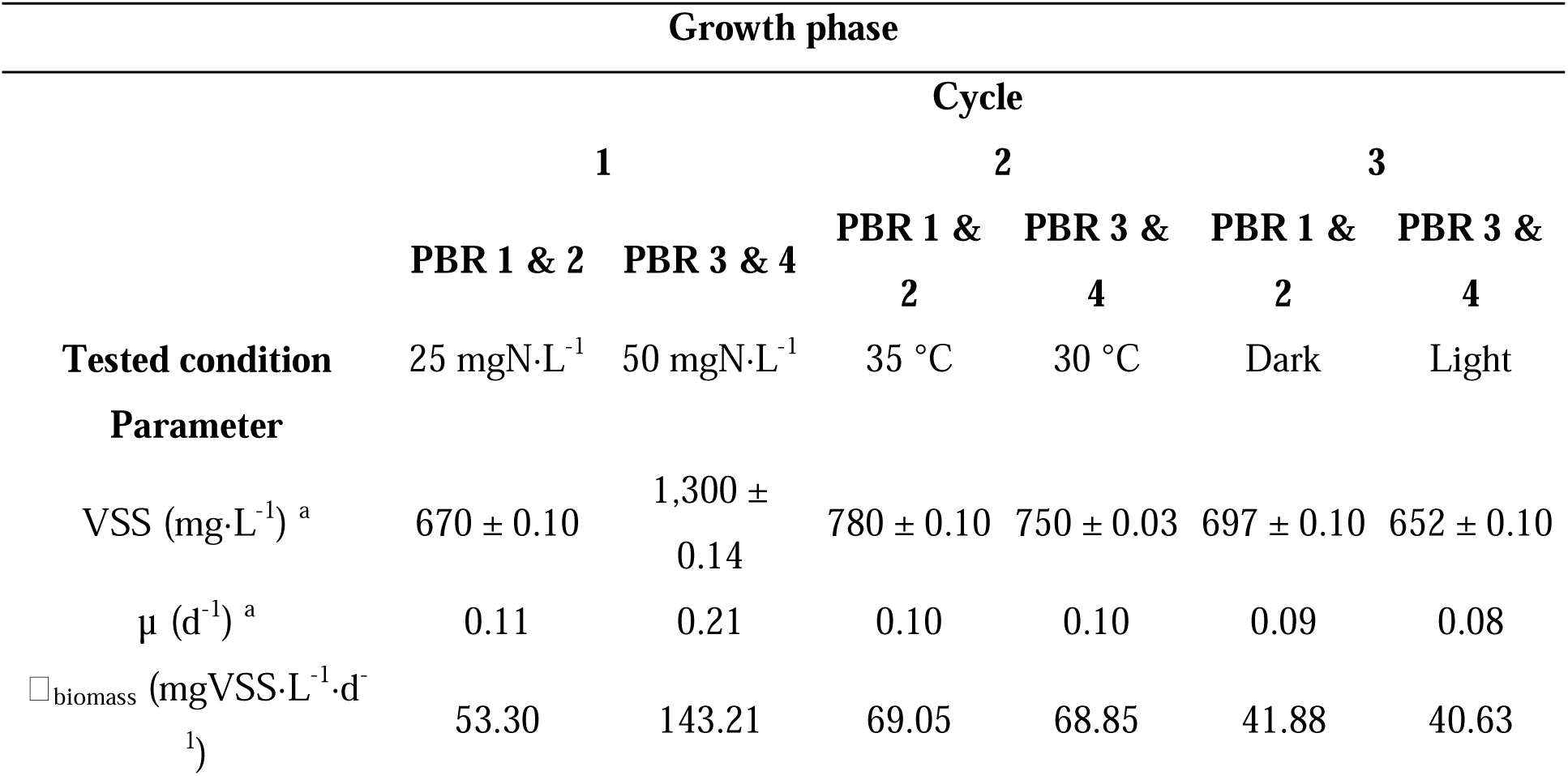
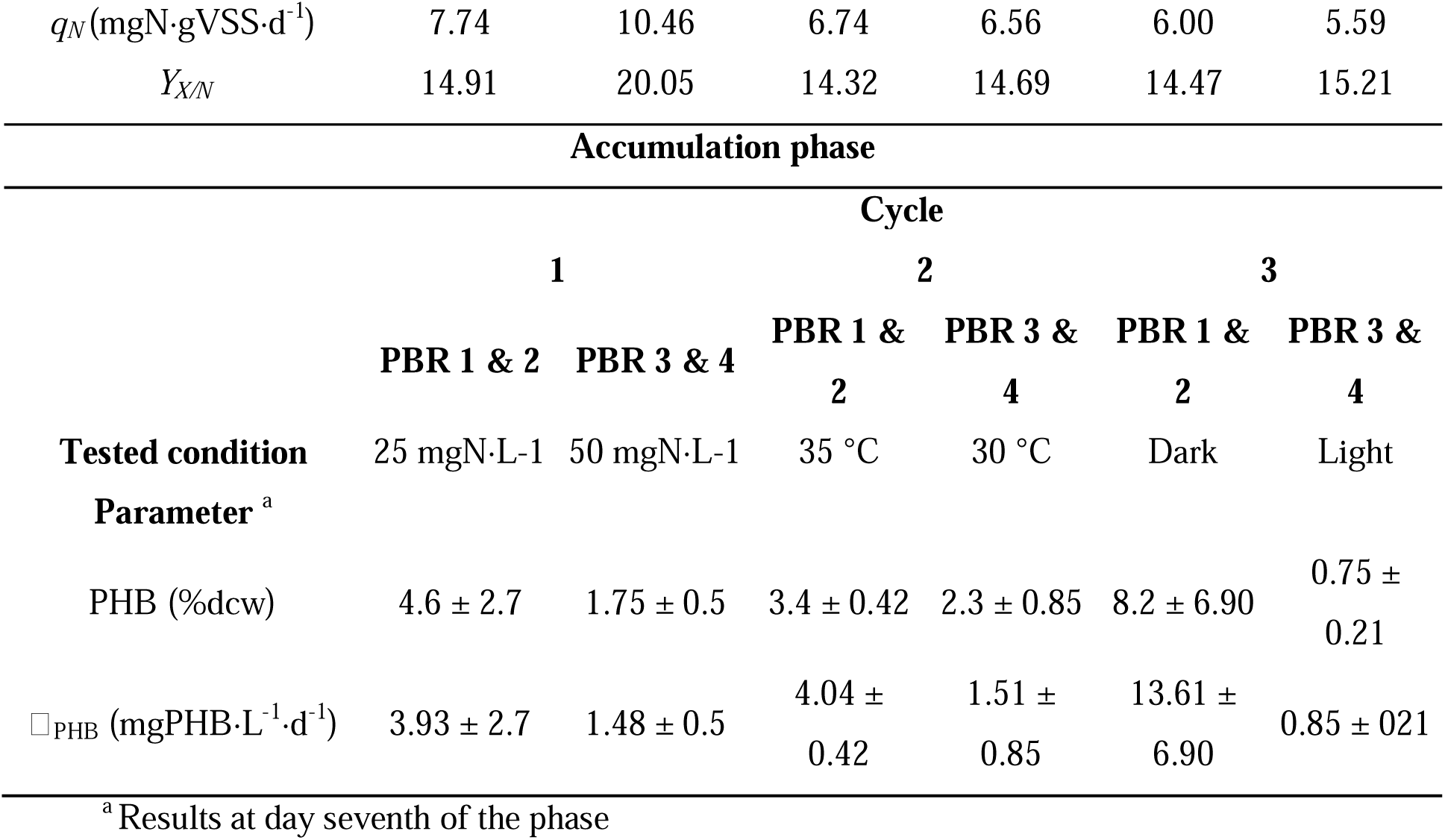
Average of the kinetic and stoichiometric parameters obtained during growth and accumulation phase of the three cycles period.

Although the initial N concentration in the growth phase of the conditioning period was equal as that settled for PBR 3 & 4 in cycle 1 (50 mg·L^-1^), the biomass grew faster during the growth phase of the conditioning period (µ = 0.31 d^-1^) than in the same phase in cycle 1 in PBR 3 & 4 (µ = 0.21 d^-1^). This difference in biomass growth could be attributed to the higher P concentration in the conditioning period (1 mg·L^-1^) and the sudden increase in green algae, which tend to grow faster than cyanobacteria (Visser et al., 2016). This would suggest that the biomass (1,120 mgVSS·L^-1^) in the conditioning period was mainly composed by green algae and in lower concentration by cyanobacteria. Moreover, it is worth noting that throughout the cycles, growth rate (µ) decreases (Table 3), while there was an increase in cyanobacteria through this period (Fig. 4A). Also, this reduction in cell growth could be attributed to the absence of external IC as bicarbonate during these growth phases, which is the primary source of inorganic carbon for autotrophic organisms in aquatic environments (Salbitani et al., 2020). While 100 mgIC·L^-1^ were added in the growth phase of the conditioning period, no IC was added during the cycles period; thus, cell relied on alternative sources of carbon (PHB) to support their metabolic needs. This could slow down the overall rate of growth (Table 3), as these stored materials may not be as readily available or efficient as bicarbonate as a source of carbon.

**Figure 4.**
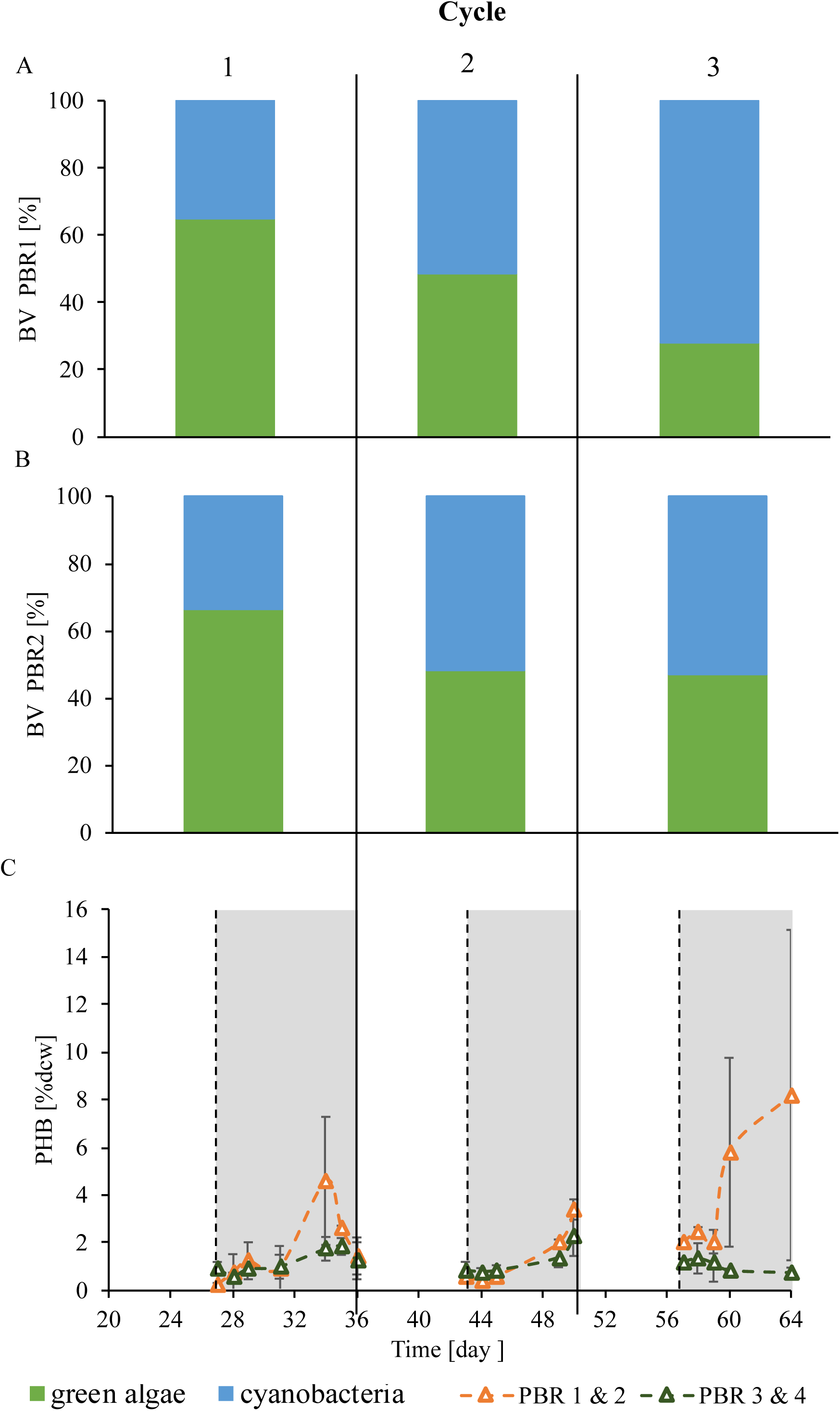
Biovolumes of (A) PBR 1 & 2 and (B) PBR 3 & 4 during the cycle period. (C) PHB evolution in the cycle period. PHB was not measured in growth phase.

The effects of temperature and light during the starvation phase were also asset in cycle 2 and cycle 3, respectively. Temperature was controlled at 35°C in PBR1 & 2 and at 30°C in PBR 3 & 4. As observed in the previous growth phase (from cycle 1), N was consumed in 7 days and biomass reached values of 780 ± 100 mgVSS·L^-1^in all the PBRs in cycle 2 and 800 ± 40 mgVSS·L^-1^ in cycle 3 (Fig. 3B). Nutrients intake also suggests that the stored PHB was being consumed. The PHB concentration decreased from the last day of starvation phase in cycle 1 to the beginning of the starvation phase in cycle 2; and from the last day of starvation phase in cycle 2 to the beginning of the starvation phase in cycle 3 were observed (Fig. 4C), meaning that the biopolymer was being consumed as a carbon source for cell growth. This observation was also supported by nitrogen intake in each growth phase (Fig. 3A and 3B).

Outcomes revealed that to promote PHB production, the best temperature was 35 °C in order to favour cyanobacterial while decreasing green algae growth, as PHB synthesis was higher in PBR 1 & 2 than in PBR 3 & 4 (Fig. 4 and Table 3). Moreover, the concentration of green algae, which had already decreased since P concentration was set at 0.1 mg·L^-1^ from cycle 1 onwards, continued to decrease until approximately 30% at the end of cycle 3 in PBR 1 & 2 (Fig.4A, 4B and A.4), demonstrating the predominance of cyanobacteria over green algae by the increase in temperature. Cyanobacteria concentration domination over green-algae by warming has also been described in (Lürling et al., 2018, 2013).

Results from cycle 3 showed that reactors should also be enclosed, meaning that no light was needed for PHB synthesis, which was in accordance with the result obtained in the Design of Experiment conducted in (Altamira-Algarra et al., 2022) with the used microbiome. The assumption that light is not required for PHB production has been proved before due to anoxic environment caused by dark cultivation conditions since photosynthetic oxygen is not produced in the dark (Koch et al., 2020; Sharma and Mallick, 2005a). This could be partially related to the increased pool of NADPH under dark (Pelroy et al., 1976), which is necessary for PHB biosynthesis (García et al., 2018; Hauf et al., 2013). Here, a notorious difference in PHB was observed between PBRs.

Production in the non-enclosed PBRs was 0.75 %dcw ± 0.21, the lowest observed in all the performed cycles (Fig. 4C and Table 3). In addition, from the 600 mgAc·L^-1^ added in the four PBRs, biomass from PBR 1& 2 consumed almost all of it (500 mgAc·L^-1^ ± 35.7), while that from PBR 3 & 4 only consumed 124 mgAc·L^-1^ ± 8.63.

### 3.3. Growth and PHB production under optimal operating conditions

Outcomes from the three cycles performed to study the effect of three ecological stresses (N concentration, temperature and light) stablished the operation mode for the next period, which was performing nine iterated cycles of biomass growth followed by PHB accumulation phase. These repetitions consisted of seven days of growing and seven days of PHB synthesis (Fig. 1).

Here two PBRs were used with the biomass from the prior PBR 1 & 2, as it was the culture with higher PHB production in the previous cycle (Table 3). Biomass increased from 400 to approximately 745 mgVSS·L^-1^ in both PBRs with the addition of 25 mgN·L^-1^ and 0.1 mgP·L^-1^ after seven days in all the repetitions (Table 4). A PHB decrease was always observed between the end of one accumulation phase (repetition *n*) and the beginning of the next accumulation phase (repetition *n*+1) (Fig. 5C); suggesting that during each growth phase, biomass used the stored PHB as carbon source, together with the external N and P to support cell growth, as already observed in the prior cycles and also by other authors (Johnson et al., 2009; Serafim et al., 2004). This trend is more clearly observed when the PHB concentrations reached at the end of the accumulation phase are somewhat low. When PHB is higher, the biopolymer concentration at the end of the growth phase is also higher, suggesting that not all PHB has been consumed.

**Tables 4.** Average and standard deviations of the kinetic and stoichiometric parameters obtained during growth of the 9 performed repetitions.

|  |  |  |
| --- | --- | --- |
| **Growth phase** |  |  |
| **Parameter** | **PBR 1** | **PBR 2** |
| VSS (mg·L<sup>-1</sup>)<sup>a</sup> | 739.78 ± 227.87 | 750.88 ± 226.57 |
| $\mu$ (d<sup>-1</sup>)<sup>*</sup> | 0.13 ± 0.04 | 0.12 ± 0.03 |
| $\square_{biomass}$ (mgVSS·L<sup>-1</sup>·d<sup>-1</sup>) | 77.78 ± 24.91 | 74.87 ± 18.42 |
| $q_N$ (mgN·gVSS·d<sup>-1</sup>) | 6.61 ± 1.86 | 6.86 ± 2.14 |
| $Y_{X/N}$ | 19.42 ± 2.37 | 18.45 ± 2.37 |

<sup>a</sup>Result at day seventh.

**Figure 5.**
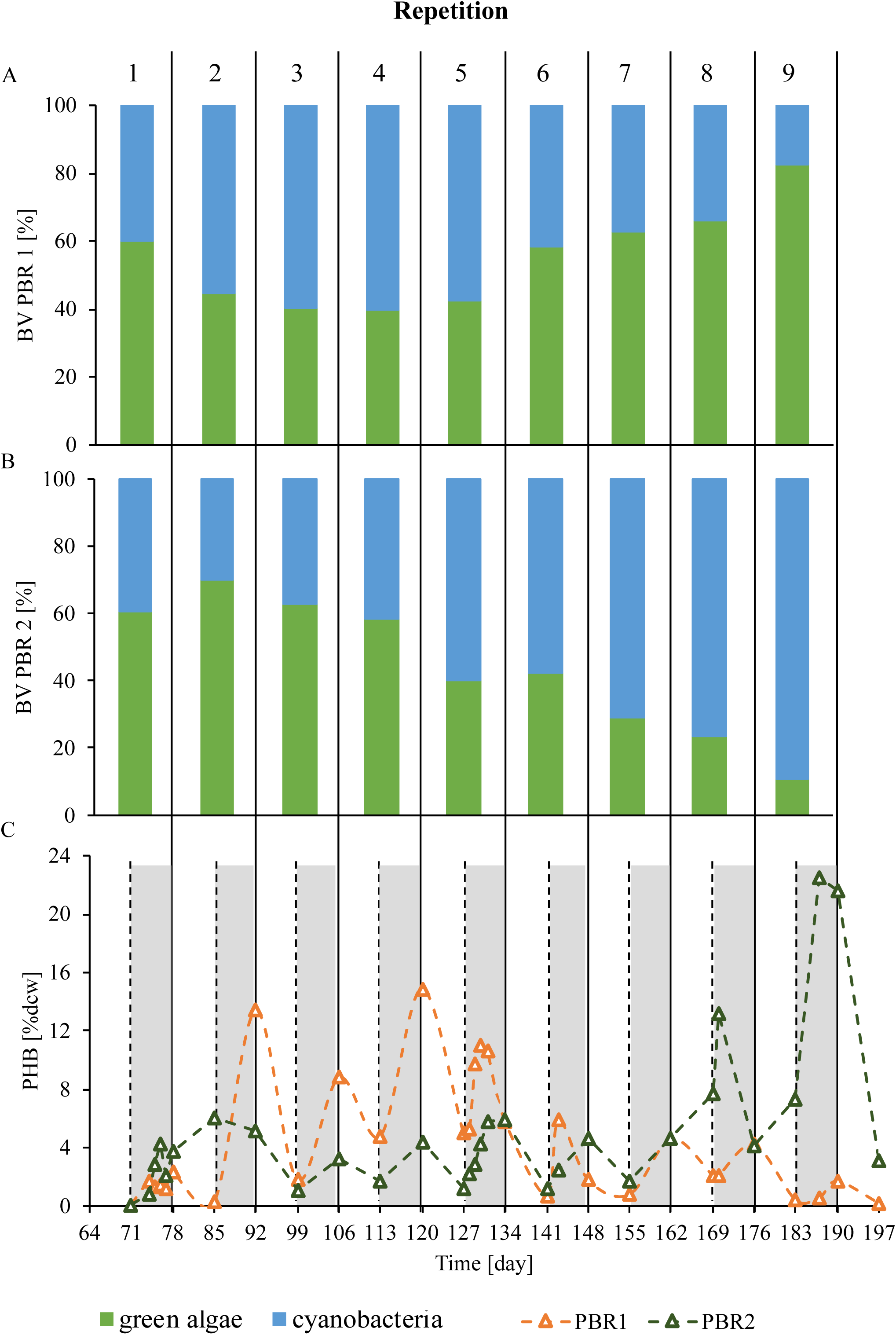
(A) Biovolume of (A) PBR1 and (B) PBR2 during the iterated period. (C) PHB evolution in the iterated period. PHB was not measured in growth phase.

In the first repetition, both PBRs had similar behaviour on PHB production (Fig. 5C). However, production by PBR 1 increased to 13.4 %dcw in the second repetition and remained in similar values at every new accumulation phase until the fifth iterated cycle (Fig. 5C). Curiously, PHB production in PBR 1 decreased to < 5 %dcw from the sixth accumulation onwards (Fig. 5C and Table 5). In PBR 2, PHB production did not underwent such a sudden rise; instead the microbiome did not produce more than 10 %dcw PHB until the repetition number eight, when PHB production was 13.2 %dcw (Fig. 5C). By the end of the last accumulation, PHB production was 22 %dcw (Fig. 5C and Table 5). Similarly, (Rueda et al., 2022a) achieved a biopolymer production rate of 26 %dcw with a monoculture of *Synechococcus* sp. with the addition of Ac after 15 days of accumulation.

**Tables 5.** Results on PHB production of each of the 9 performed repetitions. Values were calculated at day seventh of accumulation.

| Accumulation phase |  |  |  |  |  |  |
| --- | --- | --- | --- | --- | --- | --- |
| Repetition | PHB (%dcw) | | $\square_{\text{PHB}}$ (mgPHB·L<sup>-1</sup>·d<sup>-1</sup>) | | $Y_{\text{PHB/Ac}}$ (g PHB<sub>COD</sub>/g Ac<sub>COD</sub>) | |
|  | PBR 1 | PBR 2 | PBR 1 | PBR 2 | PBR 1 | PBR 2 |
| 1 | 2.3 | 3.8 | 3.14 | 5.17 | 0.13 | 0.2 |
| 2 | 13.4 | 5.2 | 8.99 | 0 | 0.24 | 0.1 |
| 3 | 8.8 | 3.3 | 4.65 | 1.46 | n.d. | n.d. |
| 4 | 14.9 | 4.4 | 10.18 | 2.72 | 0.27 | 0.09 |
| 5 | 10.6 | 5.8 | 11.98 | 9.85 | 0.24 | 0.21 |
| 6 | 1.8 | 4.6 | 1.93 | 5.96 | n.d. | n.d. |
| 7 | 4.6 | 4.6 | 5.74 | 4.38 | n.d. | n.d. |
| 8 | 4.2 | 7.7 | 3.64 | 6.24 | 0.12 | 0.36 |
| 9 | 1.7 | 21.6 | 0.88 | 9.74 | 0.04 | 0.4 |

n.d. stands for no data.

Regarding to Ac consumption, in PBR 1 half of the Ac remained in the media after seven days in the first repetition (Table A.2). Interestingly, at each new accumulation, the acetate consumed was greater, which should be attributed to an also higher biopolymer production (Table A.2). However, as mentioned, an increase in PHB did not occur (Fig. 5C), suggesting that Ac was not exclusively used for PHB synthesis. This consideration can also be seen by the Y_PHB/Ac_, which remained constant until repetition 8 and 9, when it suddenly dropped (Table 5). Contrary, in PBR 2, where cyanobacteria were more prevalent than green microalgae (Fig. 5), the concentration of Ac by the end of each repetition remained around 100 mg·L^-1^. The highest Y_PHB/Ac_ was obtained in repetition 9 (Table 5). However, the theoretical maximum Y_PHB/Ac_ that could be reached by consuming 600 mgAc·L^-1^ is 0.60, which represents 52 %dcw PHB assuming biomass concentration of 745 mgVSS·L^-1^ (the average VSS produced during the iterated cycles considering both PBRs). Here lower yields were obtained (Table 5), which suggests the possibility of optimizing Ac uptake by changing substrate addition, in order to increase PHB production and avoid an external carbon source in the coming growth phase. In this sense, one possibility could be continuous substrate feeding or pulse-wise addition. Notably, despite these differences in Ac uptake, pH during accumulation phase remained quite constant in values around 8 in both PBRs after a few days (Fig. A.5).

Interestingly, cyanobacteria proportion in PBR 1 remained quite constant until repetition 6, where it suffered a decrease from 73 % at the end of repetition 5 to 46 % at the end of the sixth repetition. By the end of repetition 9, green algae were clearly dominant in this microbiome (Fig. 5A). The reduction in Y_PHB/Ac_, due to lower PHB production and Ac uptake, seems to be linked with this decrease in cyanobacteria dominance and the increase in green algae in PBR 1 (Fig. 5). Furthermore, the microbiome composition of PBR 2 remained similar from repetition 1 to 7 (63 % cyanobacteria and 37 % green algae) (Fig. 5B). By the end of repetition 7, the % of cyanobacteria increased to 83 % and was still high for repetitions 8 and 9 (Fig. 5B), which agreed with the observed increment with biopolymer production and Y_PHB/Ac_ (Fig. 5C and Table 5).

The presented data clearly demonstrates that the microbiome’s composition, which includes cyanobacteria and green algae, led to significant variations in the PHB results. This is likely due to the fact that green algae are non-PHB-producers with a greater BV (and weight) than PHB-producers (cyanobacteria). (Fradinho et al., 2013b) tested a photosynthetic mixed culture (bacteria and algae) which produced 30 %dcw PHB due to the lower number of algae present in the culture compared to a 20 %dcw PHB production in a previous test with the same culture (Fradinho et al., 2013a). (Arias et al., 2017), who also worked with a mixed culture composed by green algae and cyanobacteria, observed that PHB was not accumulated by the lack of cyanobacteria in their cultures. The variations in PHB production (Fig. 5A) suggest that the microbiome may have produced more PHB than what was detected, which highlights the high capacity of the microbiome to produce this biopolymer.

Although the PBRs were replicas and subjected to the same operation procedures, they resulted in different outcomes. These differences arise from small deviations in macroscopic variables that are often imperceptible to the researchers. Such variations can lead to changes at the microscopic level that ultimately affect bioproduct results. For example, in our case, it is possible that slight variations in pH regulation (Fig. A5) or differences in lighting, based on the PBRs placement, could condition the evolution of the microbial populations. Therefore, meticulous control is necessary to ensure the reliability of the bioprocessing, regardless of the richness of the microbiome.

## 4. Conclusions

The experimental data demonstrated a substantial increase in PHB production from 2 %dcw PHB in the first phase to 22 %dcw PHB after a total of 179 days of reactor operation. These findings suggest the successful development of biopolymer-producing biomass. However, the presence of green algae led to a decline in PHB production, posing a challenge to maintaining consistent production rates. Although operational parameters such as P concentration or temperature could potentially control the survival of green algae, their abundance sometimes exceeds the controllable limits. This highlights the importance of implementing effective strategies to monitor and manage the growth of competing microorganisms in the reactor.

The results indicate that this strategy could offer a new methodology to enrich a photosynthetic microbiome in PHB-producers. This discovery could pave the way for more efficient PHB production strategies in the future, while also minimizing the growth of unwanted microorganisms in the reactor.

### CRediT authorship contribution statement

**Beatriz Altamira-Algarra:** Conceptualization, Validation, Formal analysis, Methodology, Investigation, Writing – original draft. **Artai Lage:** Methodology. **Joan Garcia:** Conceptualization, Resources, Writing – review & editing, Supervision, Project administration, Funding acquisition. **Eva Gonzalez-Flo:** Conceptualization, Supervision, Writing – review & editing.

## Declaration of Competing Interest

The authors declare that they have no known competing financial interests or personal relationships that could have appeared to influence the work reported in this paper.

## Acknowledgements

This research has received funding from the European Union’s Horizon 2020 research and innovation programme under the grant agreement No 101000733 (project PROMICON). B. Altamira-Algarra thanks the Agency for Management of University and Research (AGAUR) for her grant [FIAGAUR_2021]. E. Gonzalez-Flo would like to thank the European Union-Next Generation EU, Ministry of Universities and Recovery, Transformation and Resilience Plan for her research grant [2021UPF-MS-12].

## Notes

### Competing Interest Statement

The authors have declared no competing interest.

