## Supplementary material for "Species composition determines bioplastics production in photosynthetic microbiomes: strategy to enrich cyanobacteria PHB-producers"

* corresponding author

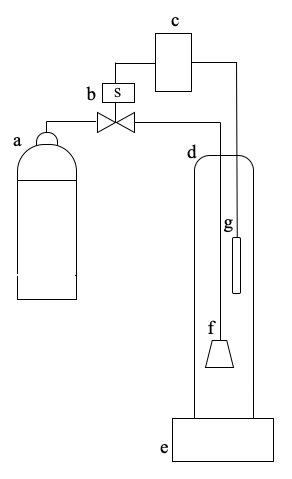

**Figure A1.** Scheme of the experimental set-up. a) CO2 bottle; b) electrovalve; c) pH controller; d) photobioreactor; e) magnetic stirrer; f) CO_2_ diffusor; g) pH probe.

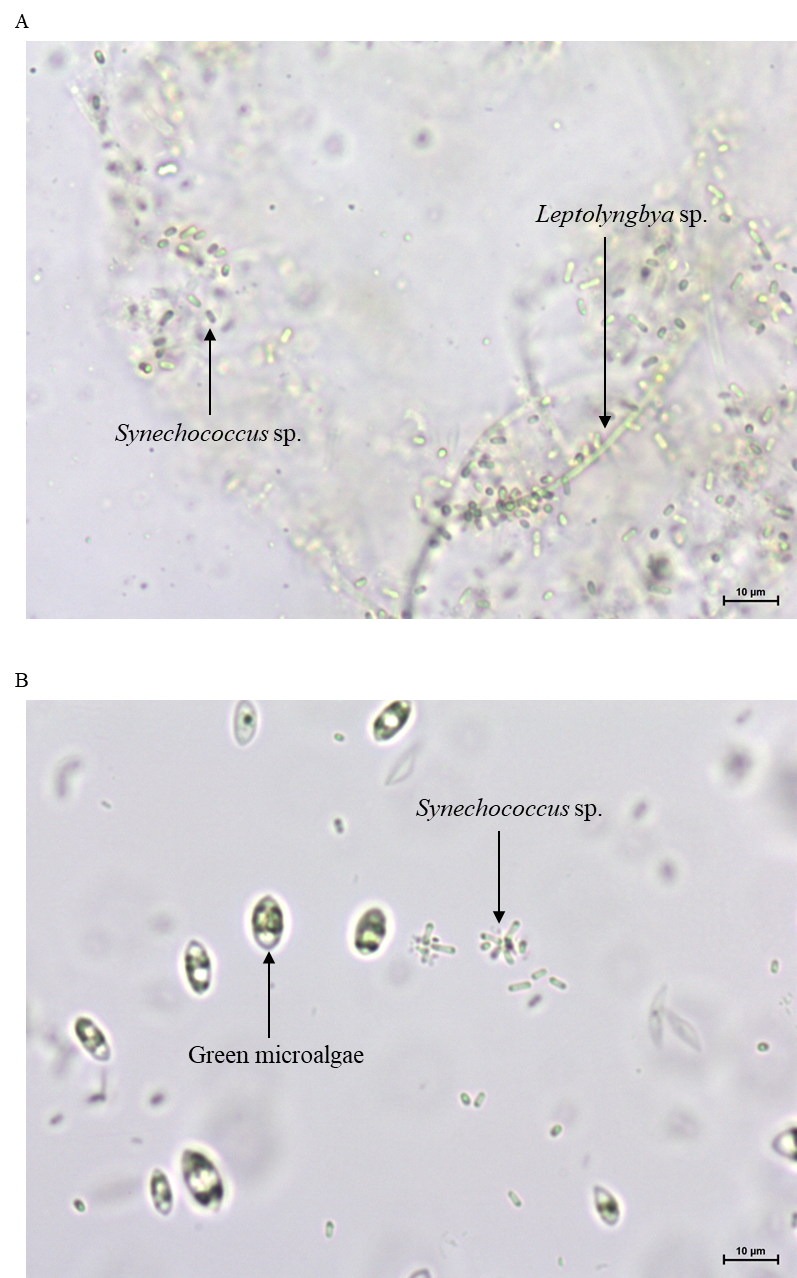

**Figure A2.** Microscope images under bright light microscopy at 40X of (A) inoculum and (B) culture at the end of conditioning period. In (A) unicellular cyanobacteria *Synechococcus* sp. and filamentous cyanobacteria *Leptolyngbya* sp. can be seen. In (B) the presence of green algae and *Synechococcus* sp. is observed.

| Turbidity [NTU] | VSS [g·L-1] |
| --- | --- |
| 218.333 | 0.344444 |
| 244.333 | 0.31 |
| 300 | 0.43 |
| 684.333 | 0.769375 |
| 749.333 | 0.8 |
| 605.333 | 0.683333 |
| 547.333 | 0.610476 |
| 402.333 | 0.630455 |
| 268.667 | 0.445 |
| 269.333 | 0.445 |
| 292.667 | 0.47 |
| 423 | 0.625062 |
| 512 | 0.67 |
| 601 | 0.759615 |
| 582.667 | 0.8 |
| 257 | 0.39 |
| 271.333 | 0.44 |
| 298 | 0.43 |
| 446 | 0.638462 |
| 513.667 | 0.691639 |
| 673.333 | 0.790833 |
|  | 0.846237 |
| 1122 | 1.36154 |
| 292.667 | 0.415 |
| 292.667 | 0.427778 |
| 299 | 0.425 |
| 366.667 | 0.573077 |
| 396 | 0.545951 |
| 498.667 | 0.580833 |
| 638 | 0.714211 |
| 758 | 1.025 |
| 239 | 0.478022 |
| 752.6 | 0.998333 |
| 280 | 0.515385 |
| 683.3 | 0.892308 |
| 806 | 1.06667 |
| 262.333 | 0.5975 |
| 593.333 | 0.85 |
| 809.333 | 1.22 |
| 360.333 | 0.547255 |
| 738.667 | 1.16115 |

**Figure A3.** Calibration curve VSS-Turbidity for microbiome CC.

**Table A1.** Data used to calculate biovolumes.

| **Cell type** | **Average width (µm)** | **Average lenght (µm)** | **Cell shape** | **Count (n)** | **Average volume (µm^3^)** | **Volume equation*** |
| --- | --- | --- | --- | --- | --- | --- |
| *Synechococcus* sp. | 1.64 | 4.29 | Cylinder | 20 | 22.72 | $v=\frac{1}{4}\pi$*l*$w^{2}$ |
| Green microalgae | 8.06 | 10.63 | Ellipsoid | 20 | 521.90 | $v=\frac{1}{6}\pi$*l*$w^{2}$ |

*In volume equation, *l* stands for average length and *w* for average width.

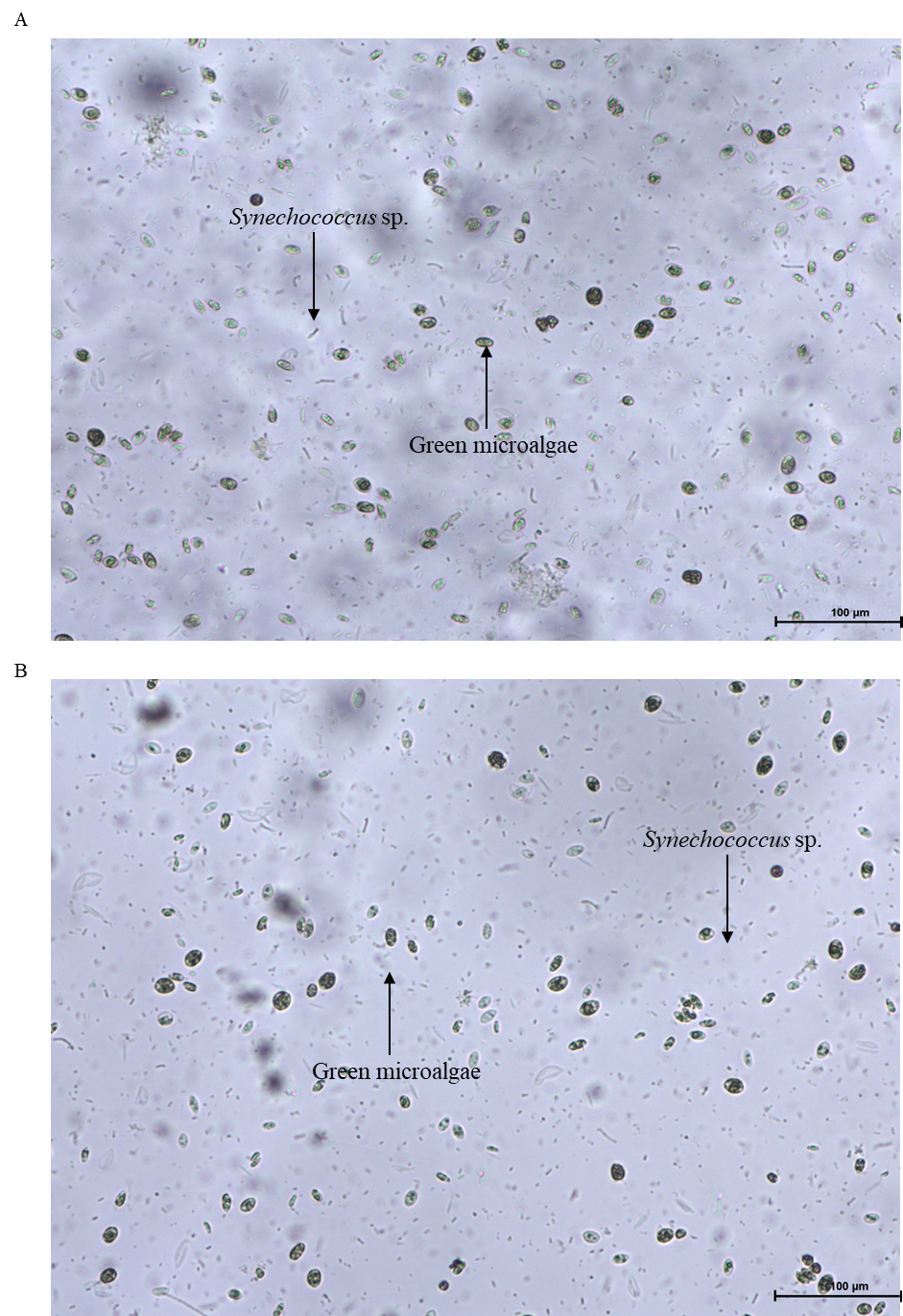

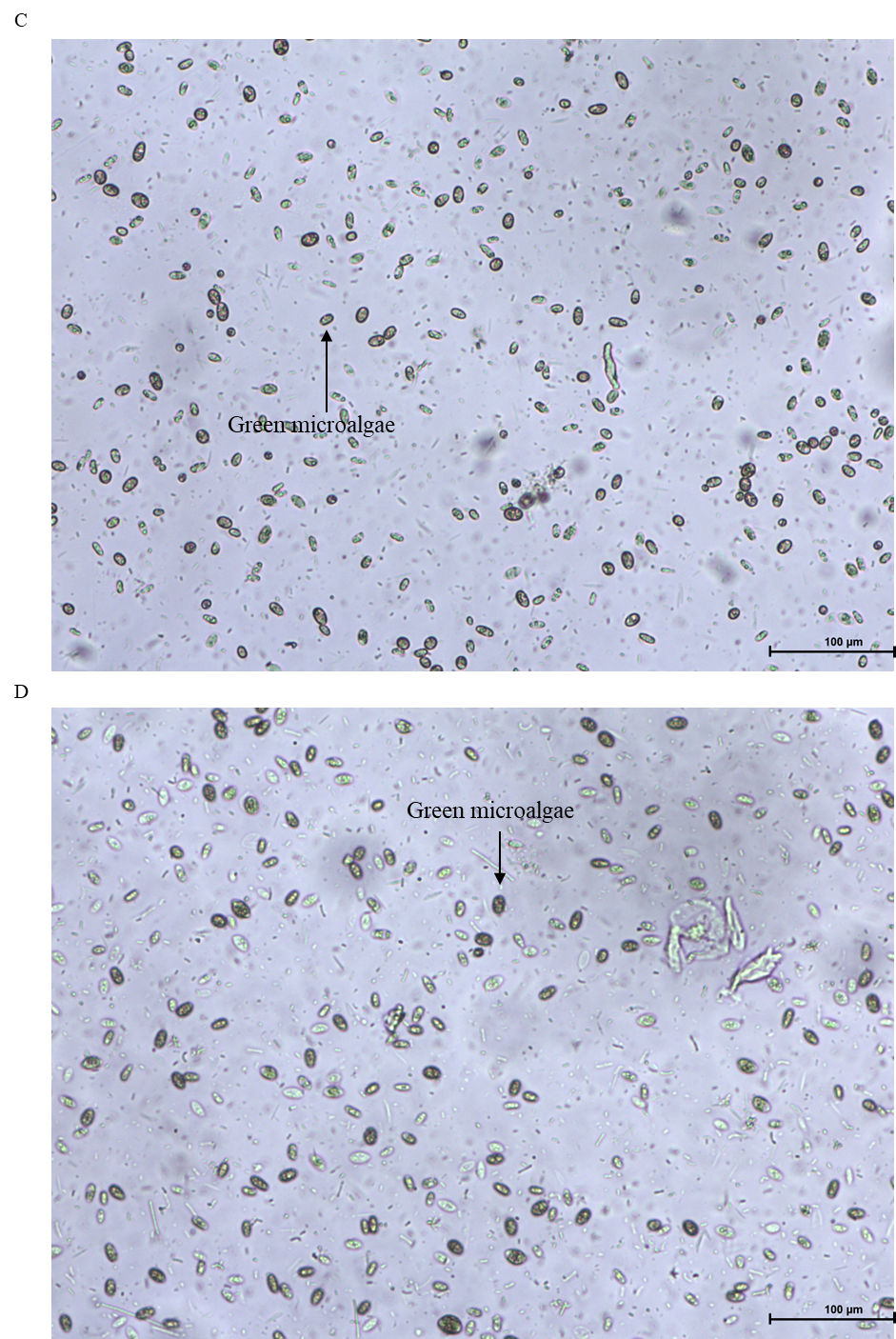

**Figure A4.** Microscope images under bright light microscopy at 10X of (A) PBR1, (B) PBR2, (C) PBR3 and (D) PBR4 at the end of cycle period. Differences in presence of unicellular cyanobacteria *Synechococcus* sp. and green algae between PBR 1 & 2 and PBR 3& 4 can be detected. In PBR 1 & 2 it can be clearly observed the lower abundance of green algae.

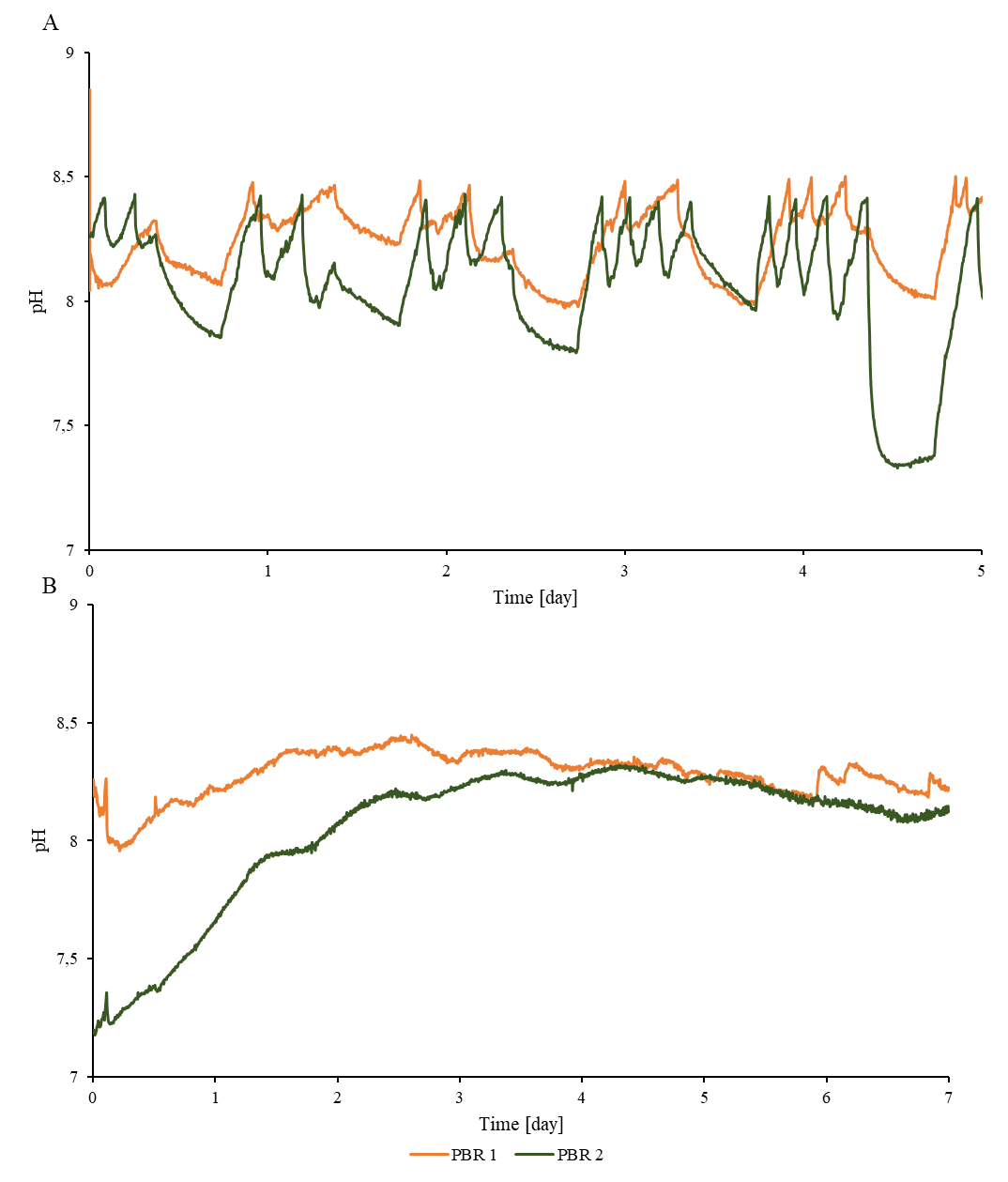

**Figure A5.** Example of pH evolution through on cycle. (A) Profile during growth phase in repetition 5 and (B) during accumulation phase in repetition 4.

**Table A2.** Consumed acetate during the iterated period. Values were calculated by subtracting the amount of Ac remaining in the PBRs to the initial Ac concentration (600 mg·L^-1^).

| **Repetition** | **Consumed Ac (mgAc·L^-1^)** | |
| --- | --- | --- |
|  | **PBR1** | **PBR2** |
| 1 | 277.62 | 292.079 |
| 2 | 424.71 | 454.46 |
| 3 | n.d. | n.d. |
| 4 | 596.94 | 523.824 |
| 5 | 600 | 342.06 |
| 6 | n.d. | n.d. |
| 7 | n.d. | n.d. |
| 8 | 218.169 | 449.3 |
| 9 | 296.73 | 444.58 |
